# Phylogeny-based selection of representative variants with Navargator: Proof of principle using humoral cross-reactivity data from two immunization studies

**DOI:** 10.1101/2024.09.08.611914

**Authors:** David M. Curran, Dixon Ng, John Parkinson, Trevor F. Moraes, Scott D. Gray-Owen, Jamie E. Fegan

## Abstract

**Background:** The accessibility to the immune system of bacterial surface proteins makes them attractive targets for subunit vaccines. However, this same property also means they tend to exhibit high sequence variability. Achieving broad cross-protection usually necessitates that antigens from multiple isolates are included, but the choice of sequence variant is a non-trivial problem. Visual inspection of phylogenetic trees is the norm, but this is subjective and can be greatly influenced by the choice of viewing software. This has real-world implications, as groups have shown that the selection of non-optimal antigens likely led to lower cross-protection in the commercially available vaccines against *Neisseria meningitidis* serogroup B.

**Aim / Methods:** To address this problem, we have developed Navargator, bioinformatics software that takes a phylogenetic tree as input and identifies the variants that are the most similar to the greatest number of other sequences. The underlying premise is that cross-reactivity will be correlated with phylogenetic distances extracted from the tree; this was validated by several rodent immunization studies with the proteins transferrin-binding protein B and factor H binding protein from *N. meningitidis* and *N. gonorrhoeae*, measuring antibody-based cross-reactivity between an antigen panel using a custom high-throughput ELISA.

**Results:** Navargator has been made freely available both as an online tool and as source code for local installation. We implemented several different clustering methods, with exact algorithms for smaller datasets, and heuristics suitable for large trees of thousands of sequences. Our immunization studies have shown that this approach is sound, and that cross-reactivity is predicted well by phylogenetic distances in a sigmoidal manner.

**Conclusions:** The complexity of vaccine development rises sharply with each additional antigen included, so using the minimal number required is an important consideration. Navargator attempts to facilitate this in a systematic and generalizable manner. The user can run the analysis by selecting their desired number of representatives, or they can provide any form of cross-reactivity data and have the program identify a minimum reactivity threshold via correlation with the phylogenetic tree. The program will then identify the smallest number of representatives required to satisfy this threshold.

## Introduction

As sequencing technology improves in both read length and accuracy and the cost of performing sequencing has decreased, the quantity of sequence information available in public databases has increased exponentially over the last two decades. This is particularly true of sequence information related to pathogens that are highly monitored, both as part of routine surveillance as well via clinical diagnostics, while also frequently undergoing high levels of mutation due to evolutionary pressure asserted by antibiotics, vaccines, and the immune response. This was exemplified by the explosion of sequence information collected over the nearly five years of the COVID-19 pandemic, where nearly nine million nucleotide and forty-six million protein sequences of SARS-CoV-2 genomes have now been deposited in the NCBI Virus database (https://www.ncbi.nlm.nih.gov/labs/virus/vssi/#/virus?SeqType_s=Nucleotide&VirusLineage_ss=taxid:2697049).

While newly emerging pathogens such as SARS-CoV-2 develop new variants which may outcompete previous variants, endemic pathogens often have numerous strains circulating concurrently, which offers challenges when developing vaccines to target these pathogens. *Streptococcus pneumoniae*, an important cause of respiratory and invasive infections in young and old populations, represents an extreme example in that it exists as 100 different serotypes (Ganaie et al., 2020), defined by the polysaccharide capsule surrounding the cell, each of which is serologically distinct. Because of this, current vaccines include either twenty (PCV-20) to twenty-three (PPSV-23) different antigens. Protein subunit vaccines are an attractive alternative yet offer a different challenge since protein variants are more mutable than the repetitive subunits in polysaccharide capsules; this is particularly true with surface-exposed proteins, which frequently undergo high levels of antigenic variation in an attempt to subvert host immunity.

Phylogenetic analyses are useful to understand the sequence diversity in a population, and performing clustering on a phylogenetic tree partitions that population into groups that share similarities. This is particularly useful is in the selection of antigens for novel recombinant protein vaccines. An ideal vaccine will be allele-agnostic with respect to the target; that is, it will protect against strains expressing all possible forms of the target antigen. However, the ability of a vaccine to elicit broad cross-protection is tied to population-level properties of the antigen. One of the most important is the level of sequence variation found in populations of the infectious agent. If a strain possesses a target protein that is very similar to the antigen used in a vaccine, then it is very likely that the immune response generated by the vaccine will react to that strain. Conversely, an immune response is more likely to fail as the difference increases between the target protein and the vaccine antigen. To protect against a population of pathogens with high levels of sequence variation of the target, one approach is to formulate a vaccine with multiple variants of the antigen, as is done with the 3-valent polio vaccines, 4-valent dengue virus vaccine (Dengvaxia, Sanofi Pasteur) or 9-valent human papillomavirus vaccine, Gardasil-9 (Merck).

However, increasing the number of variants used in a vaccine increases the difficulty of production and licensing, so the ideal vaccine will use the fewest variants necessary to achieve the desired level of cross-protection.

While it may seem conceptually simple to look at a phylogenetic tree and identify a variant that is nearby to many neighbours – akin to glancing at a scatterplot and picking one point near the middle of a group of points – this is a common fallacy, and the reality is that a tree is a very different form of representing information. On a 2D scatterplot, both spatial dimensions represent some form of distance (regardless of the units). On the common rectangular format of displaying a phylogenetic tree, only the dimension stretching from tips to root represents distance. The other spatial dimension does not represent any form of distance; it conveys some information about branching patterns, but mostly exists for the sake of legibility. Thus, the most appropriate method is to measure the total branch distance between each variant, and then select that which is most proximal to all other variants. Since this is not feasible when large numbers of sequence variants are present, we have developed and here describe a novel software and graphical user interface (GUI) that allows for rapid, user-driven clustering of phylogenetic data and the identification of central variants using a variety of methods. Towards validating the utility and validity of this approach, we have evaluated the cross-reactivity of the neisserial transferrin binding protein B (TbpB), an antigenically variable surface-anchored lipoprotein that binds host transferrin and aids in iron acquisition during infection, and which has been proposed as a desirable vaccine target against both pathogens in the genus, *Neisseria meningitidis* and *Neisseria gonorrhoeae*. We then use Navargator to consider the variability among naturally occurring factor H binding protein (fHbp) sequences expressed by *N. meningitidis*, and to estimate how representative the fHbp that are present in two approved serogroup B meningococcal vaccines, Bexsero and Trumenba, and then test this prediction with immune cross-reactivity studies.

## Results

The pathogenic *Neisseria* include two species are important to human health; *Neisseria meningitidis*, the causative agent of invasive meningococcal disease, and *Neisseria gonorrhoeae*, the cause of the second most prevalent bacterial sexually transmitted disease, gonorrhea. Surface antigens expressed by these bacteria are highly variable due to their phase and antigenic variability, as well as their natural competence for genetic transformation by environmental DNA (Rotman & Seifert, 2014). *N. meningitidis* is encapsulated and, as such, has been primarily targeted with highly efficacious conjugate capsular vaccines (Tzeng & Stephens, 2021). However, the capsule of serogroup B *N. meningitidis* is non-immunogenic in humans, and thus significant effort has gone towards the development of protein-based vaccines to target strains within this serogroup. Indeed, there are now two licensed vaccines for prevention of serogroup B (MenB) strains (Findlow et al., 2020): Bexsero, composed of an outer membrane vesicle derived from the bacteria and three recombinant proteins as target antigens, *Neisseria* adhesin A (NadA), neisserial heparin binding protein (NHBP), and factor H binding protein (fHbp); and Trumenba, composed of a bivalent formulation of two fHbp variants from each of the two phylogenetically-defined subfamilies. *N. gonorrhoeae* lacks a licensed vaccine for prevention, however, presumably due to it sharing many surface antigens with *N. meningitidis*, epidemiological studies have suggested some cross protection for individuals immunized with Bexsero (Abara et al., 2022; Ladhani et al., 2024; Petousis-Harris & Radcliff, 2019).

### Selection of immunogens and ELISA panels

Due to its essential function for both pathogenic *Neisseria* species, we have experience targeting their transferrin receptor as a vaccine target. This receptor is comprised of an integral outer membrane protein (transferrin binding protein A, TbpA) and a lipid-anchored transferrin binding protein B (TbpB), which is highly exposed on the bacterial surface. To analyze the natural variation among TbpBs from these two species, we generated a phylogenetic tree depicting their sequence diversity (Figure 1A).

**Figure 1.**
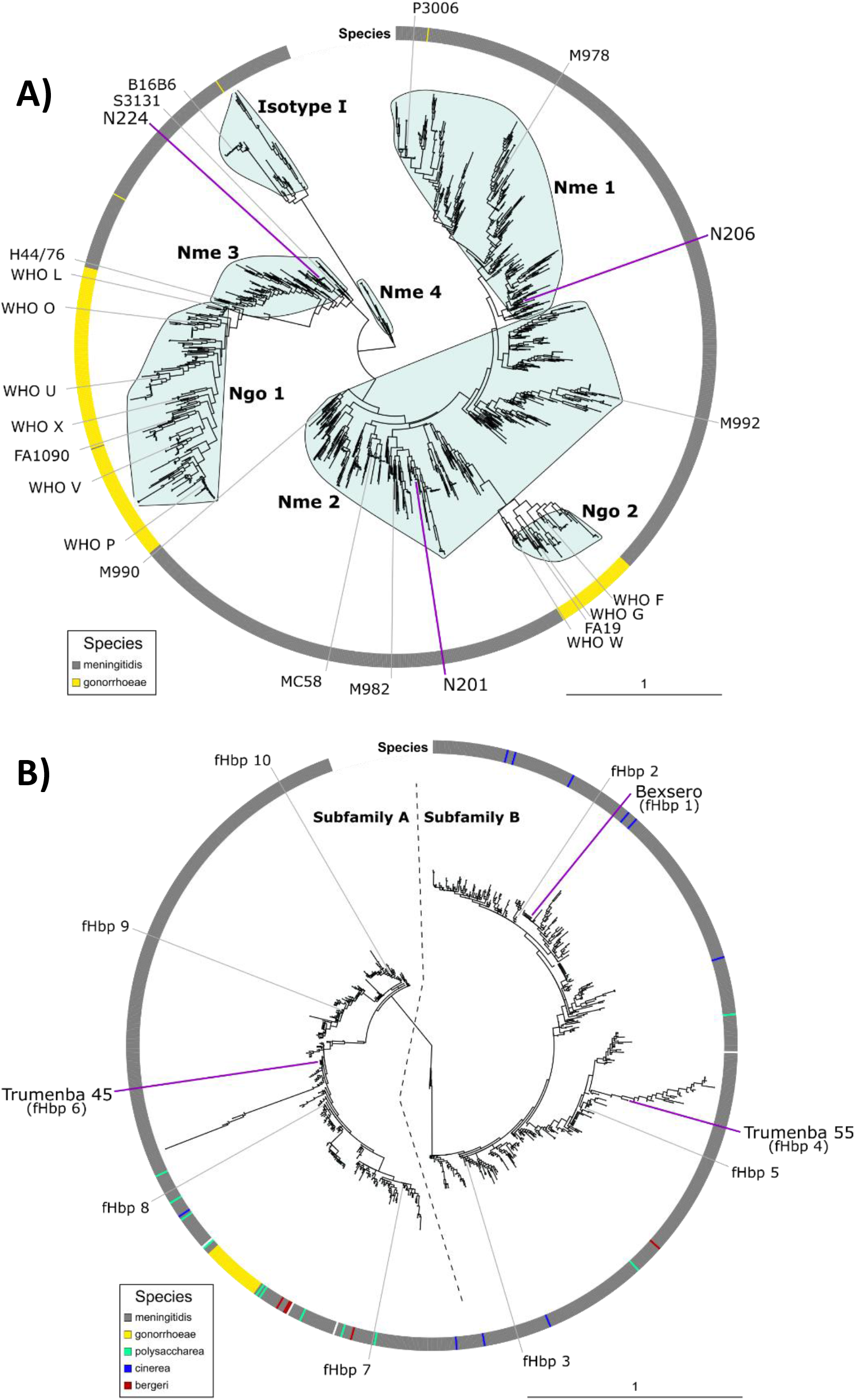
Phylogenetic trees showing the diversity between A) TbpBs from *N. meningitidis* and *N. gonorrhoeae*, and B) fHbps from various species of *Neisseria*. The ring around each tree indicates the originating species for each sequence. The positions of the antigens used in in our immunizations are indicated on the trees with purple lines, while the additional variants used in our ELISA panels are indicated with grey lines. Both trees are organized into subgroups to facilitate discussion, with A) showing 7 clusters identified by Navargator, and B) showing the classification used in the literature.

From this, we used Navargator to consider different clustering patterns, settling on 7 as this was the smallest pattern that separated groups in agreement with existing observations, such as separating the Isotype I from the rest and separating *N. gonorrhoeae* sequences from *N. meningitidis*. We then selected one immunogen from each of the three major *N. meningitidis* clusters Nme1, Nme2, and Nme 3. These were not selected to be the most central or potentially cross-reactive variants – as one might do when designing a final formulation – but because we had experience formulating them in vaccines and wanted to quantify cross-reactivity across a wide range of distances. Similarly, the ELISA panel was chosen not because they were equally spaced around the tree, but because this was a reasonably-sized subset of commonly used strains with variable representation.

We wanted to examine the cross-reactivity elicited by a second class of antigen, and so we decided to select as our immunogens the existing human vaccines targeting fHbp: Bexsero and Trumenba. To select our ELISA panel, we generated a phylogenetic tree depicting the diversity of publicly available fHbp protein sequences (Figure 1B). Navargator was used to select 10 representative variants from this tree, constrained so that three of those variants were the antigens in the two vaccines. The fHbp proteins were cloned into our custom vector that tags them with a streptavidin-binding peptide (Fegan et al., 2021) and then expressed in *E. coli*. Nine of these were able to effectively bind human Factor H after being expressed (the exception being fHbp 5), indicating they were properly folded upon capture to the neutravidin coated ELISA plate (data not shown), and so constitute our ELISA panel.

### TbpB reactivity is correlated with phylogenetic distance

To test the patterns of cross-reactivity between the TbpBs, we synthesized the chosen genes, expressed them as recombinant proteins to immunize mice, and then functionally tested cross-reactivity via an ELISA panel consisting of 23 *Neisseria* TbpB proteins. The results indicate high reactivity against the homologous (immunizing) TbpB variants, as expected (Figure 2A) for N201, N206, and N224 immunogens. When evaluating cross-reactivity – as measured by optical density of the colorimetric reaction from the ELISA – there was generally high levels of cross-reactivity against TbpBs from the same or nearby clusters, however the reactivity dropped substantially when tested against the Isotype I TbpB (B16B6), as well as when tested against any gonococcal TbpBs. When correlating the cross-reactivity with phylogenetic distance from the immunizing TbpB, we again saw high reactivity against closely related TbpBs and minimal reactivity with highly distant TbpBs (generally, a phylogenetic distance greater than 3) (Figure 2B). TbpBs falling in intermediate phylogenetic distances from the immunizing antigen showed high variability in their reactivity. This may be due in part to immunodominant epitopes on the immunizing antigen, but also results from the normal effect of variability in immune responses between individual animals, a kind of phenomenon referred to as a “batch effect”. This can be seen most prominently in the N206 graph, where the data from individual animals forms consistent striated layers, and the individually fit curves spread apart from the collective curve. For example, at the intermediate distance sera from “N206 2” reacts much stronger than sera from “N206 1”, while for the gonococcal panel members the reactivity pattern follows “N206 1” > “N206 3” > “N206 2” > “N206 4”.

**Figure 2.**
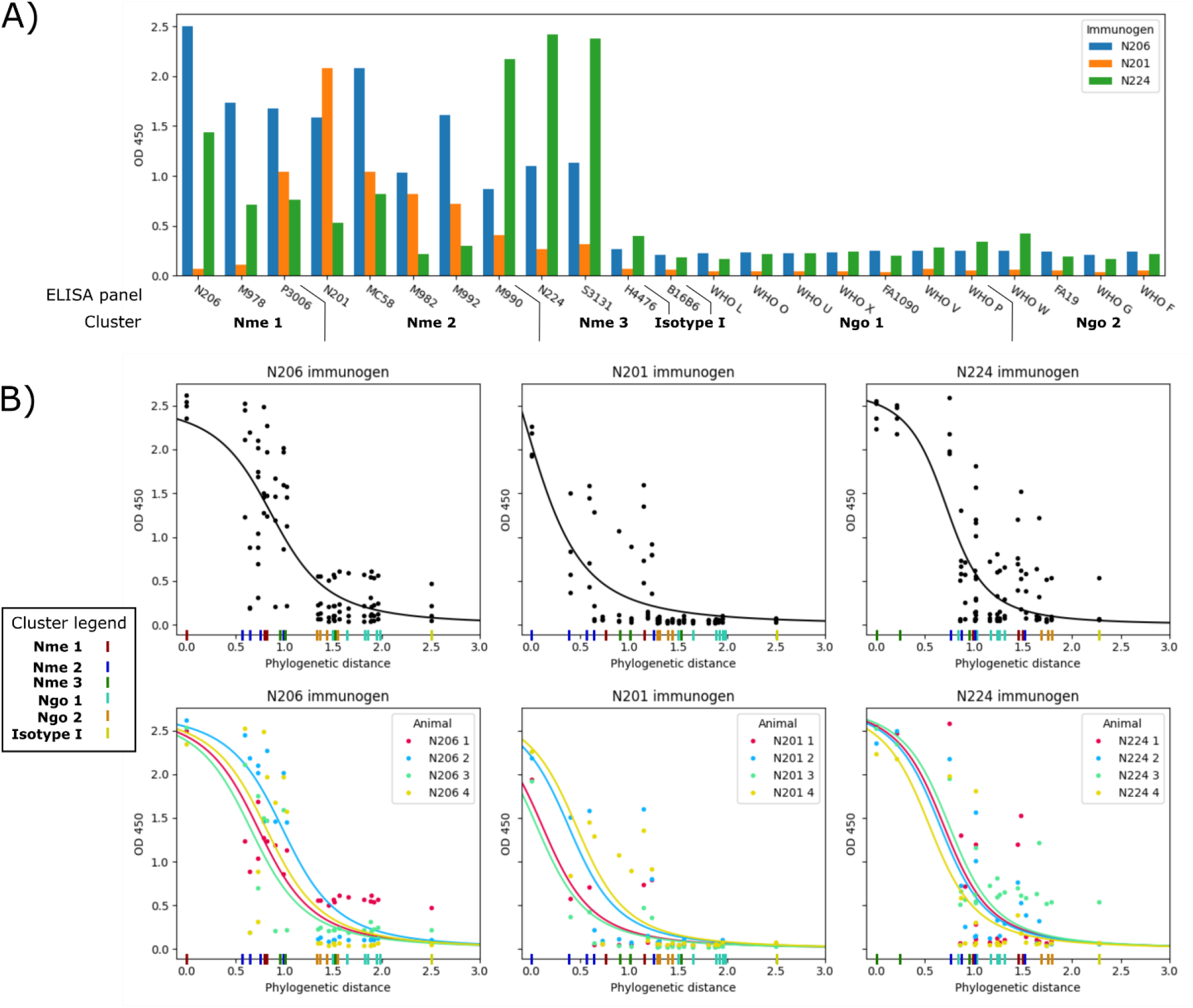
Immunogenicity and cross-reactivity of the three TbpB immunogens. A) shows how reactive each immunogen is to our ELISA panel and to which cluster (Figure 1A) that panel member belongs, while B) shows how that reactivity relates to the phylogenetic distance between immunogen and ELISA panel member. Each column of graphs represents one immunogen where both graphs display the same set of points. The top graph in each column shows the curve fit to all of the data, while the bottom shows curves fit to each individual animal. The coloured dash marks on the x-axes indicate which cluster that member of the ELISA panel is from.

### fHbp reactivity is correlated with phylogenetic distance

Once establishing the utility of Navargator in selecting antigen variants to formulate a vaccine, we aimed to consider the validity of using Navargator to understand the breadth of cross-reactivity offered by two different vaccines already licensed for use in humans. While *N. meningitidis* and *N. gonorrhoeae* both express TbpB (Figure 1A), fHbp is generally only expressed by *N. meningitidis* (Figure 1B). Both vaccines for serogroup B meningococci contain fHbp as a recombinant antigen. Important for binding to the host complement inhibitor Factor H, variants of fHbp are separated into two subfamilies, subfamily A and subfamily B (Fletcher et al., 2004), which share approximately 60-75% sequence homology at the nucleotide level. Bexsero, which contains only one variant of fHbp (fHbp 1, subfamily B), likely relies on the synergistic effect of the multiple components that its comprised of to elicit broad cross-protection (reviewed in (Viviani et al., 2022)), while Trumenba incorporates one lipidated fHbp variant from each subfamily to elicit broad coverage (fHbp 6 from subfamily A, Pfizer ID fHbp-45; and fHbp 4 from subfamily B, Pfizer ID fHbp-55) (Jiang et al., 2010). Previous work has interrogated the selected variants, finding that immunization with sequence variants more central to the subfamily elicited serum that was more broadly bactericidal to strains across the subfamily than do the variants that are included in either MenB vaccine (Konar et al., 2013). This finding makes it enticing to consider whether the robust use of phylogenetic data could select more broadly cross-protective vaccines.

To test this, we evaluated whether the breadth of fHbp-induced cross-reactivity correlated with the phylogenetic distance between the included fHbp variant and members of the ELISA panel. Mice were immunized with a 1/5^th^ human dose of either Bexsero – containing fHbp 1 from Subfamily B – or Trumenba – containing fHbp 6 from Subfamily A and fHbp 4 from Subfamily B. After two doses of the respective MenB vaccines, mouse serum was tested against the fHbp ELISA panel (Figure 3A). As expected, the highest reactivity was against the fHbp variant used for immunization, with immune sera from mice receiving Bexsero reacting higher to fHbp 1 compared to sera from mice that received Trumenba, which reacted higher to fHbp4 and fHbp6. Notably, Trumenba sera showed a strong bias towards the fHbp 4 antigen, suggesting that this variant may be immunodominant in mice. This is particularly evident when correlating the reactivity against phylogenetic distance. When the distance was measured to the fHbp 4 antigen (Figure 3B, middle column) the expected relationship was observed, with cross-reactivity decreasing as phylogenetic distance increased. However, when measuring the distance to the fHbp 6 antigen (Figure 3B, right column), the relationship was inverted.

**Figure 3.**
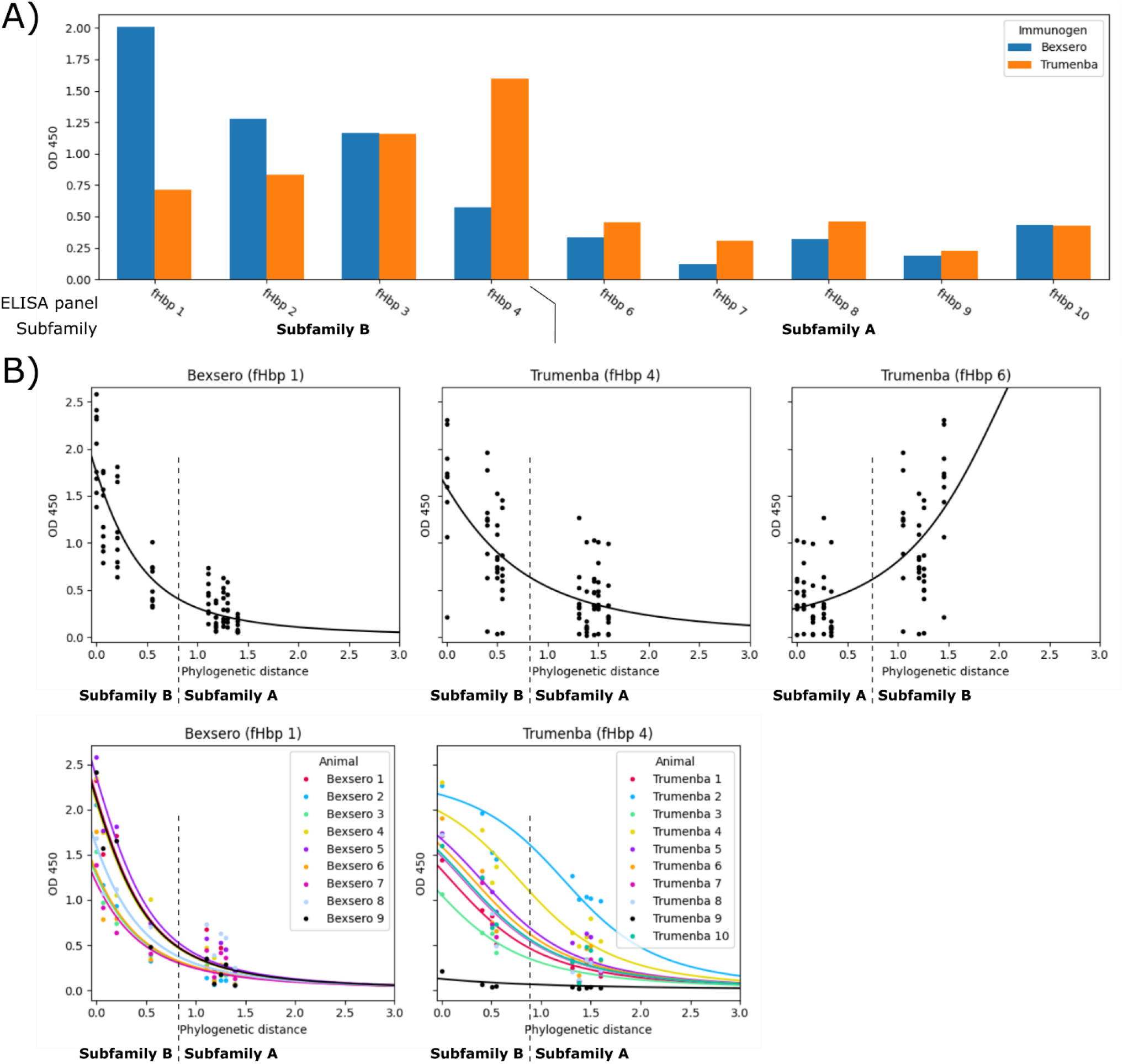
Immunogenicity and cross-reactivity of the two MenB vaccines. A) shows how reactive each vaccine is reactive to our ELISA panel and to which subfamily (Figure 1B) that panel member belongs, while B) shows how that reactivity relates to the phylogenetic distance between immunogen and ELISA panel member. Each column of graphs represents one immunogen where both graphs display the same set of points. The top graph in each column shows the curve fit to all of the data, while the bottom shows curves fit to each individual animal. As Trumenba contains two fHbp antigens, the middle column measures the distance from fHbp 4 to the ELISA panel, while the right column measures from fHbp 6.

Similar to the TbpB results (Figure 2), some batch effects were seen in the Bexsero-induced immune sera responses, but they were very strong in the Trumenba sera (Figure 3B, bottom middle graph). Here we see a strong hierarchy of reactivity, with sera from animal “Trumenba 1” always responding the strongest followed by “Trumenba 4”, and sera from “Trumenba 9” showing virtually no activity to any member of the ELISA panel.

## Methods

### Reconstructing the TbpB and fHbp phylogenies

All TbpB sequences from *N. meningitidis* or *N. gonorrhoeae* were downloaded from the PubMLST database (Jolley et al., 2018), supplemented by 60 sequences from our lab collection. The signal peptides were removed using SignalP v5.0 (Almagro Armenteros et al., 2019), as well as the anchor peptides (approximately the first 20 residues of the mature peptide). All duplicate sequences were filtered leaving 1485 unique TbpBs, 1178 from *N. meningitidis* and 307 from *N. gonorrhoeae*.

All sequences annotated as “fHbp” or “fHbp_peptide” from all species of *Neisseria* were downloaded from the PubMLST database, and signal peptides were removed using SignalP v5.0. All duplicate sequences were filtered leaving 898 unique fHbps from the species *N. meningitidis* (838), *N. gonorrhoeae* (30), *N. polysaccharea* (11), *N. cinerea* (10), *N. bergeri* (5), *N. uirgultaei* (1), *N. maigaei* (1), *N. blantyrii* (1), and *N. subflava* (1).

Alignments were generated using MAFFT v7.475 (Katoh & Standley, 2013) running the E-INS-i algorithm. Phylogenetic trees were generated using RaXML v8.2.12 (Stamatakis, 2014) using the PROTGAMMAWAG model of evolution.

### Cloning and expression of the immunogens and ELISA panel

Nucleotide sequences were codon optimized for expression in *E. coli* and synthesized on a template vector (Integrated DNA Technologies); the synthesized sequences contain flanking regions of complementarity to the downstream expression vector and could be amplified and subcloned by restriction-free cloning techniques (van den Ent & Löwe, 2006). All primers were synthesized by Sigma Aldrich, and Phusion DNA polymerase (Thermo Fisher Scientific) was used in PCR reactions under manufacturer recommended conditions. Amplification product from the template vectors generated megaprimers that were purified by gel electrophoresis and excised following manufacturer instructions (Bio Basic). Purified megaprimer fragments were used in a secondary linear amplification reaction around the expression vector template. The reaction was incubated with 1 unit of *Dpn*I (Thermo Fisher Scientific) at 37°C for 6 hours to degrade the methylated template. 8 μL of the reaction mixture was used to transform chemically competent *E. coli* MM294 cells, followed by 1 hour of recovery in Luria Broth (LB) at 37°C shaking. Transformants were selected by plating on LB agar plates supplemented with ampicillin (100 μg/mL). Single colonies were used to inoculate 5 mL of LB cultures and incubated overnight at 37°C shaking. Plasmids from each clone were extracted using a MiniPrep kit following manufacturers recommendations (Bio Basic) and the insertion was confirmed by Sanger-sequencing (The Center for Applied Genomics, TCAG).

Protein production was carried out in *E. coli* T7 express cells (New England Biolabs) which were made chemical-competent and transformed with expression vectors by heat-shock. Cultures recovered in LB broth for 1 hour at 37°C with shaking at 175 RPM, before selection by adding 3 mL of LB supplemented with ampicillin (100 μg/mL) and grown for another 3 hours. This starter culture was then used to inoculate 6 L of autoinduction media (Studier, 2014) with 50 μg/mL of ampicillin, which was then shaken overnight at 37°C at 175 RPM for 16 hours and a further 24 hours at 20°C. Cells were centrifuged at 5000 x g’s for 30 minutes at 4°C, resuspended in 50 mL of resuspension buffer (50 mM Tris pH 8.0, 300 mM NaCl) supplemented with protease inhibitors (cOmpleteTM Mini, Roche), 1 mg/mL lysozyme, and 0.03 mg/mL DNase I. Cells were homogenized (EmulsiFlex-C3, Avestin) and the resulting lysate was clarified by centriguation at 16,000 x g’s for 90 minutes at 4°C, followed by 0.22 μm syringe filtration (Millipore). The clarified lysate was then recirculated on a HisTrap EXCEL FF column (Cyutiva) overnight which is then mounted onto an ÄKTA Purifier 100 (GE Healthcare), washed and eluted with imidazole (50 mM Tris pH 8.0, 300 mM NaCl, 300 mM imidazole pH 7.4). The eluted fraction was dialyzed into exchange buffer (50 mM Tris pH 8.0, 150 mM NaCl) and in-house prepared TEV protease was added to remove the N-terminal maltose binding protein fusion partner. The cleaved sample was then dialyzed into a low salt buffer (50 mM Tris pH 8.0, 10 mM NaCl) and further downstream purification was carried out by anion exchange on a HiTrapQ FF column (Cytiva) and size exclusion on a HiPrep 26/60 Sephacryl S-200 HR (Cytiva). A final polishing step by a strong anion exchange MonoQ 5/50 GL column (Cytiva) removes lipopolysacchrides and other contaminants, before the samples were concentrated and flash frozen by liquid nitrogen.

### Collection of sera

All animal work was performed under animal use protocol 2001139, approved by the Animal Care Committee at the University of Toronto. Mice were maintained under specific pathogen free conditions and were provided water and rodent chow ad libitum.

Identified TbpB proteins were formulated into single antigen vaccines (10 µg per dose) in formulation with aluminum hydroxide gel (100 *μ*g per dose, Alhydrogel, Sigma Aldrich) in sterile phosphate buffered saline (PBS). C57Bl/6 female mice (4-5 weeks old at time of purchase, Charles River) were acclimated to the animal facility for one week, randomized into groups of 4, and then vaccinated with 100 *μ*l of the relevant TbpB vaccine or Alhydrogel only on day 0, 21, and 42. Post dose 1 and 2 serum samples were collected via saphenous bleed on days 20 and 41, while terminal serum samples were collected two weeks after the third vaccine dose via cardiac puncture after CO_2_ euthanasia. Serum samples were stored at -20 °C until use.

To generate anti-fHbp sera, C57Bl/6 male and female mice (4-5 weeks old at time of purchase, Charles River) were acclimated to the animal facility for one week, randomized into groups, and then vaccinated with either Bexsero (4CMenB, GSK, lot 9KC24) or Trumenba (bivalent rLP2086, Pfizer, lot FH4329) (n=5 male and n=5 female mice per vaccine). Vaccines were diluted 1:2 in sterile PBS and immunized intra-peritoneally at a 1:5 human dose (200 *μ*l of a 50% diluted vaccine). Mice received doses on day 0 and day 21 and on day 35 mice were humanely euthanized via CO_2_ overdose and serum was collected via cardiac puncture. Serum was stored at -20°C until use.

### ELISA cross-reactivity analysis

TbpB and fHbp ELISAs were performed as previously described (Fegan et al., 2021). Briefly, TbpBs and fHbps of interest were cloned into custom expression vectors containing his-bio-mbp-tev-tbpB constructs and transformed into *Escherichia coli* strain ER2566. Transformed colonies were grown overnight in 20 mL of ZYP-5052 autoinduction media at 37 °C with shaking. The following day, crude lysate was collected by pelleting the culture and resuspending in a 1:10 volume of lysis buffer (50mM NaH_2_PO_4_, 300 mM NaCl, 10mM imidazole, pH 8.0). 1 mM glass beads were added to the cell suspension and cells were lysed via shaking with the cell disruptor. Homogenate was centrifuged for 20 min at 4 °C to pellet cell wall material. Crude cell lysate was further diluted 1:5 in PBS and then 20 *μ*l per well was added to 384 well plates that were previously coated with 10 *μ*g/ml neutravidin (NeutrAvidin biotin-binding protein, Thermo Fisher Scientific) overnight, washed with PBS containing 0.05% tween 20 (PBST), and blocked with 5% bovine serum albumin (BSA, BioShop Canada). For TbpB evaluation, protein lysate was incubated for 2 hours at room temperature, washed, and then serum samples diluted in 1% BSA and 0.2 mg/ml MBP were added to wells at a dilution of 1:1,000, 1:10,000 and 1:50,000, and human transferrin (hTf) conjugated to HRP was used as a control to evaluate TbpB folding on the plate. For fHbp evaluation against the homologous fHbp variants, serum was added in two-fold dilutions from 1:250 up to 1:2,048,000, and the endpoint titre was considered the last dilution in which a signal above background (mean plus three standard deviations). For fHbp evaluation against the ELISA panel, serum was diluted in 1% BSA and added to wells at a dilution of 1:50,000 and 1:250,000, with serum from transgenic mice expressing human Factor H added at a 1:100 dilution as a fHbp folding control. Serum was incubated overnight at 4°C, plates were washed, and secondary antibody (goat anti-mouse IgG H&L (HRP), 1:10,000 dilution, Abcam, cat. ab6789) was added for 2 hours at room temperature. Plates were washed and developed with KPL SureBlue TMB Microwell Peroxidase Substrate (SeraCare, cat. 5120-0077). Reactions were quenched with 2 N H_2_SO_4_ and read at 450/570nm.

### Clustering overview

Classification of variants, chosen / available / ignored / unassigned Setup of variables and definitions used in following sections:

Generally speaking, a clustering algorithm operates on some dataset *U* = {*u*_1_, …, *u*_*n*_} consisting of *n* members, and outputs a partition of these data into *k* subsets *C* = {*c*_1_, …, *c*_*k*_} such that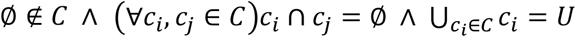 each subset is non-empty, mutually disjoint, and their union covers the whole dataset. Each of these subsets defines one cluster and taken together the set *C* is referred to as the clustering pattern.

Navargator performs clustering operations on the set *L* = {*l*_1_, …, *l*_*n*_} consisting of *n* leaves extracted from some phylogenetic tree *T*, where *D*(*i, j*) gives the phylogenetic distance between leaves *i* and *j*. For our application we are more concerned with the representative variant from each cluster as opposed to the clustering pattern itself, so the output of all of our algorithms is the set of medoids *M*, consisting of *k* leaves, each chosen as the representative for one cluster. We can define the function 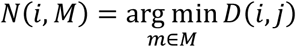 (written simply as *N*(*i*) when *M* is obvious from context) that returns the nearest medoid to a given leaf, and the function *H*(*m*) = {*l* ∈ *L*|*N*(*l*) = *m*} that returns the set of leaves for which *m* is the nearest medoid. Then the clustering pattern as defined above can be generated by assigning each leaf to the cluster represented by the nearest medoid *C* = {*H*(*m*)|*m* ∈ *M*}. As we are concerned with the phylogenetic distance between the leaves of *T* we evaluate clustering patterns using a tree score function *S*(*M*) = ∑_*i*∈*L*_ *D*(*i, N*(*i, M*)) that sums the distance between every leaf and its nearest medoid.

### k-medoids clustering

The first class of clustering algorithm has the number of clusters *k* specified by the user, and has the useful property that no information about the relatedness between data needs to be specified beforehand.

#### Brute force

This algorithm is only suitable for small to medium sized trees if *k* is very small (in practice no larger than 3 or 4). The algorithm simply evaluates all possible combinations of medoid choice, and returns the selection with the lowest tree score. There are 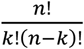 possible ways to choose *k* medoids out of *n* choices, which is why the runtime increases exponentially as *k* or *n* gets larger.

#### k-medoids heuristic

We implement a version of the k-medoids algorithm (Kaufman & Rousseeuw, 1987). We also implement a permutation of that algorithm, CLARA, designed to handle large datasets (Gentle et al., 1991).

### Radial threshold clustering

One class of clustering algorithm seeks to partition nodes into the minimum number of clusters such that the maximum distance between any pair of nodes in a cluster is less than or equal to some threshold *t*. This is referred to as a “maximum diameter” objective function, and it has the useful property that the number of clusters does not need to be specified beforehand. QT clustering (Heyer et al., 1999) is an early application of this form of algorithm to the task of identifying genes with similar expression profiles. As our focus is on selecting cluster medoids, we use a related “maximum radius” objective function such that the maximum distance between the medoid and any node in the cluster is less than or equal to some threshold *t*.

Navargator implements two forms of radial threshold clustering, a greedy solver suitable for large trees, and a slower optimal solver suitable for small and medium trees. The greedy solver is just an implementation of QT clustering and proceeds as follows:

From the set of all unclustered leaves, identify the leaf with the greatest number of neighbors within distance *t*. Ties are broken by the smallest sum of distances from the leaf to its neighbors. Add this leaf to the set of medoids *M*, and remove it and all of its neighbors from the unclustered leaves. Repeat until there are no unclustered leaves remaining.

We made an interesting observation when designing an exact solver for radial threshold clustering: the uncomplicated case – where all leaves are classified as available – is an instance of the well-known minimal dominating set problem from graph theory (Haynes et al., 2013). To show this, consider a graph *G* = (*V, E*) where *V* = {*v*_1_, …, *v*_*n*_} is a set of vertices corresponding to the leaves from *T*, and *E* = {{*u, v*}|*u, v* ∈ *V* ∧ *u* ≠ *v* ∧ *D*_*uv*_ ≤ *t*} is a set of edges connecting all of the vertices for which their corresponding leaves in *T* are within the threshold distance *t*. A dominating set *D* of *G* is defined as *D* ⊆ *V*, ∀*u*(*u* ∈ *V* \ *D*) ∃*v*({*u, v*} ∈ *E*) a subset of vertices such that every vertex is in *D*, or has a neighbour in *D*; a dominating set is minimal if it has the smallest possible size. Relating this back to tree clustering, a minimal dominating set defines the smallest set of cluster medoids, such that every leaf in *T* is no further than distance *t* from at least one medoid. This is exactly the definition of radial threshold clustering.

One large improvement was to partly collapse maximal cliques in *G*. Specifically, those vertices with identical closed neighbourhoods. The closed neighbourhood of a vertex *N*[*v*] is the subgraph of *G* induced by *v* and all vertices adjacent to *v*. For two vertices *u* and *v* to have identical closed neighbourhoods, they must be connected to exactly the same set of vertices, which means they are interchangeable when identifying a minimal dominating set.

From the definition of our radial threshold problem, if the distance between leaves *u* and *v* is greater than *t*, then they will never be found in the same cluster. We extend this simple concept to the observation that if some branch in the tree is longer than *t*, then the leaves on one side of the branch will never be found in a cluster with leaves on the other side. This allows us to conceptually split *T* into a set of smaller subtrees, where our algorithm only needs to consider the leaves of one subtree at a time.

In terms of the graph *G*, this is akin to splitting it into a set of components; the minimal dominating set of *G* is equal to the union of the minimal dominating sets of the components of *G*. This dramatically lowers the complexity of the problem, and due to the polynomial nature of this algorithm in practice this translates to reducing the runtime by up to 10,000x (from 20 minutes to 50 milliseconds).

### Phylogenetic distance correlations

A sigmoidal equation was developed to model the relationship between ELISA reactivity and phylogenetic distance: 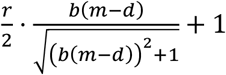, where *d* is the phylogenetic distance. The parameters optimized to fit the observed data are *b* (the slope of the curve), *m* (the distance where the response is at 50% of maximal), and *r* (the maximal response). Curves of this form are constrained to have a minimal response at 0. The ‘curve_fit‘ function from the Python package Scipy (Virtanen et al., 2020) was used to perform the curve fitting, and we found the performance was best when using the parameter bound *r* ≤ 5 which was a practical consideration as the machine we used to measure ELISA OD could not return readings above this value.

## Discussion

Here, we present novel software, Navargator, that utilizes phylogenetic distance to aid in the selection of central variants from a requested number of clusters from any phylogenetic tree towards aiding in superior antigen selection for inclusion in vaccines. We have validated that approach by comparing phylogenetic distance with cross-reactivity of immune serum from mice immunized with either a pre-clinical TbpB-containing vaccine and with anti-fHbp cross-reactivity from two human licensed MenB vaccines, Bexsero and Trumenba. Navargator is freely available both as a web tool (https://www.compsysbio.org/navargator) and as an installable package (https://github.com/dave-the-scientist/navargator) and aims to aid in the rational selection of protein immunogens for more effective, broadly protective vaccines, as well as to consider the relative potential of cross-reactivity with existing vaccines.

The problem of variant selection for inclusion in protein-based vaccines has long been underappreciated in the field of vaccinology. Variant selection has often been based on highly virulent strains, which is of high importance to protect against but may not lead to broad protection, or based on what strains are commonly utilized in a laboratory and thus have more pre-clinical information on proteins in those strains, such as is the case with *Glaesserella parasuis* and the Nagasaki strain (Hau et al., 2021). When available sequence information is utilized, the choice of variant is informed by phylogenetic analysis but the selected variant is frequently chosen by eye, an approach that is particularly challenging as phylogenetic trees become larger and complex branching patterns obscure the true relationships.

This software has proven useful to our group analyzing many varied projects – including engineering loopless antigens (Nik’s LCL paper, submitted), developing a sufficiently reactive vaccine with a minimum number of antigens (Jamie’s bivalent paper, submitted), identifying the functionality of different PmSLP variants (Islam et al., 2023), and classifying putative heme-binding proteins (Shin et al., 2024). Our software provides two major forms of clustering to the user, depending on whether the number of clusters, or whether the variation contained within each cluster, is more important for the user’s application.

Our focus here has been towards developing superior protein based vaccines, but phylogenetic trees do not have to be based on sequence data; Darwin’s original tree was based on phenotypic traits (Darwin, 1859), and morphological phylogenies were the norm until molecular phylogenies became common in the genomics era. Even more generally, a tree is just a data structure representing the relationships between a set of objects, and they appear in diverse fields of study like structural biology (protein structural similarities), linguistics (shared phonemes between languages), and computer science (file compression activity). Navargator makes no assumptions about the underlying data of a tree, and so can cluster any form of bifurcating tree. This extends its use cases far outside of vaccinology, or even biology in general.

